# Label-free imaging of 3D pluripotent stem cell differentiation dynamics on chip

**DOI:** 10.1101/2022.08.30.505835

**Authors:** Scott Atwell, Dominik J. E. Waibel, Sayedali Shetab Boushehri, Carsten Marr, Matthias Meier

## Abstract

The dynamic chemical and architectural microenvironments of 3D stem cell cultures can be controlled by integration into a microfluidic chip. Massive parallelized 3D stem cell cultures for engineering *in vitro* human cell types require new imaging methods with high time and spatial resolution to fully exploit technological advances in cell culture. Here, we introduce a label-free deep learning method called Bright2Nuc to predict *in silico* nuclear staining in 3D from bright-field images obtained using traditional confocal microscopy. Bright2Nuc was trained and applied to several hundred 3D human induced pluripotent stem cell cultures differentiating towards definitive endoderm on a microfluidic platform. Combined with existing image analysis tools, Bright2Nuc segmented individual nuclei from bright-field images, quantified their morphological properties, predicted stem cell differentiation state, and tracked the cells over time. Our methods are available in an open-source pipeline that enables researchers to upscale 3D cell phenotyping in stem cell culture.

## Introduction

New cell culture technologies for pluripotent stem cells are central to enabling *in vitro* disease models, regenerative therapies, and animal replacing drug screens ^1^. 3D culture techniques have become commonly used in differentiation trials of stem cells towards defined cell types, such as beta cells ^2^, hepatocytes ^3^, and intestinal epithelial cells ^4^. Although 3D cultures can mimic the architecture of the tissue niche, promoting cell-to-cell and cell-to-matrix interactions, the chemical microenvironment requires additional technological tools. Microfluidic chip technologies can fill this gap by automating fluid programs to test complex differentiation protocols in parallelized stem cell cultures ^5^. Further, microfluidics offers miniaturized solutions for unifying the shape and size of 3D stem cell cultures, but also for positioning the cell cultures to simplify high-content data acquisition. With the progression of cell culture technologies, analytical methods to phenotype 3D cell cultures under higher throughput conditions have become difficult because cells in 3D adopt high-density compact configurations, and cell morphologies are more heterogeneous and less recognizable than in 2D cultures. Thus, tracking and phenotyping massively parallelized 3D cultures at the single-cell level presents the challenge of not only developing image acquisition methods allowing to resolve the 3D cell cultures in high time and spatial resolution, but also powerful computational methods to handle the increasing amount of data being generated ^6^.

Owing to the widely accessible equipment, confocal fluorescence is the gold standard for obtaining single-cell resolutions in optical microscopy. Real-time imaging of cell types, functions, or cell states is hampered by the need for fluorescent reporter cell lines, which often require lengthy processes to engineer and validate ^7^. Furthermore, fluorescent reporters may affect biochemical phenomena or cell types of interest and induce cytotoxic stress under prolonged imaging times. Label-free microscopy, based on deep learning image processing and analysis, has emerged as an alternative to fluorescent reporters. For example, *in silico* staining can be used to predict fluorescent markers from the bright-field images of various tissue types ^8–11^. Once trained, deep-learning models are fast and consistent in their predictions. For high-content imaging in screening studies, label-free microscopy can dramatically reduce the image acquisition time by inferring multiple fluorescent *in silico* stains from single images without being limited by spectral cross-talk. Most of the previous approaches have focused on high-resolution imaging of 2D cell cultures or tissue sections with high numerical apertures. Higher-throughput imaging results in a trade-off between acquisition time, resolution, and phototoxicity. In particular, a higher resolution is only achieved with liquid immersion objectives, which are cumbersome for high-content data acquisition. Further, though not unprecedented ^12^, applications of *in silico* staining to live 3D cell cultures are rare. Finally, no previous approach has addressed the problem of phenotyping stem cells in a 3D environment from bright-field images.

Herein, we present a microfluidic platform for complex fluid programming in a multiplexed high-throughput manner to enable the differentiation of human induced pluripotent stem cells (hiPSC) in 3D, while simultaneously facilitating live imaging. We focused on acquiring imaging data on 3D cell cultures with a relatively low-resolution and low-numerical-aperture air-immersion 20X objective through confocal microscopy to increase data acquisition throughput. We introduced a deep learning-based method, named Bright2Nuc, to predict nuclear fluorescence within on-chip 3D cell cultures from low-magnification bright-field images. *In silico* staining images were used to infer the differentiation state of human stem cells and cell dynamics within on-chip cultivated 3D cell cultures undergoing endodermal differentiation. For reproducibility and applicability, the Bright2Nuc deep learning method and associated data were published using open-source software.

## Results

### Imaging of 3D human stem cell cultures on chip

We developed a microfluidic large-scale integration chip platform to automate the formation, culture, and differentiation of 128 3D cell cultures under 32 independent chemical conditions (**Fig. 1a**). Cells were seeded as a single-cell suspension in each chamber, and after 4 h, 3D cell cultures were formed by self-aggregation (**Fig. 1b**). A chamber height of 50 µm constrained the 3D cell cultures between the glass substrate and PDMS channel top, ensuring imaging from top to bottom (**Fig. 1c**). Together with the position stability of the cell culture compartments, the chip enabled high-content time-resolved imaging of reproducibly homogeneous (**Supplementary Fig. 1a**) live 3D cell cultures with temporal control of chemical conditions.

**Figure 1.**
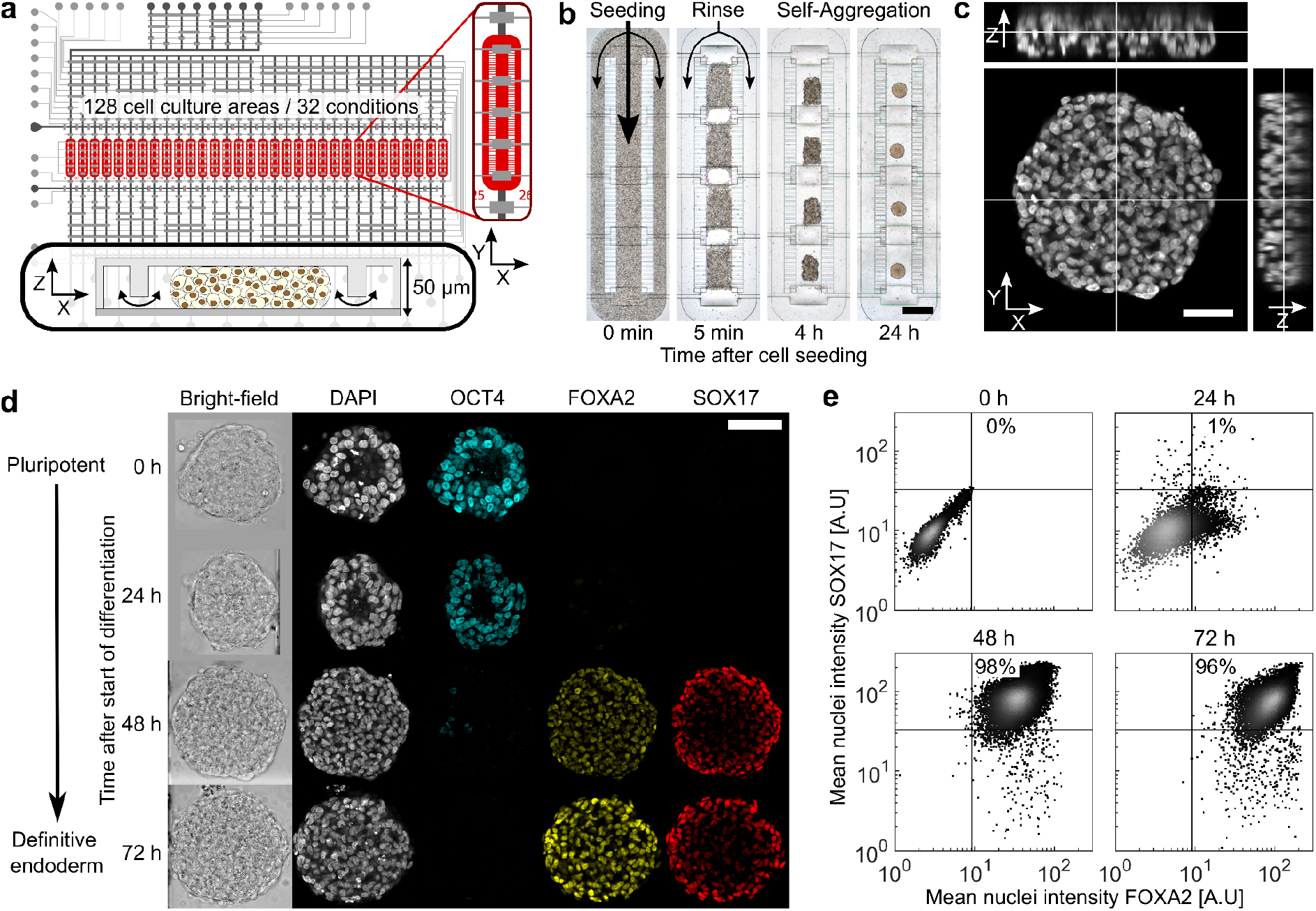
Microfluidic large-scale integration chip platform chip enables the multiplexed differentiation and imaging of 3D human induced pluripotent stem cell cultures. **a**, Scheme of microfluidic large-scale integration chip layout. The black and gray lines denote the flow and control microchannels of the double-layered PDMS chip, respectively. The inset on the right shows a magnified view of one cell culture chamber (red). The bottom inset shows the cross-section through a filled cell culture area. **b**, On-chip 3D cell culture formation process: i) seeding of single-cell solution, ii) separation of cell culture areas upon actuation of pneumatic membrane valves, followed by cell rinsing, and iii) self-aggregation of cells in the confined cell culture volume after 4 h and 24 h. Scale bar: 500 µm. **c**, Orthogonal views of an on-chip 3D cell culture stained with DAPI. Images were taken with a 0.25 µm/px xy-resolution and 1 µm/px z-resolution. Scale bar: 50 µm. **d**, Immunofluorescence confocal images of 3D cell cultures fixed along the differentiation trajectory from the pluripotency (OCT4 positive) to the definitive endoderm (FOXA2 and SOX17 double-positive) cell stage. Scale bar: 100 µm. **e**, Quantitative image analysis of cell type marker expression along the differentiation trajectory within all cells of 3D cell cultures from a single chip run. Each data point represents the mean fluorescence intensity of individual nuclei segmented in 3D; the indicated percentages represent the fraction of FOXA2/SOX17 double-positive nuclei (nuclei considered: *n*_0h_ = 5501, *n*_24h_ = 8700, *n*_48h_ = 22587, *n*_72h_ = 14936) from multiple 3D cell cultures (*N*_0h_ = 12, *N*_24h_ = 25, *N*_48h_ = 24, *N*_72h_ = 30).

Within the first 24 h after the start of DE differentiation, 3D cell cultures lost an average 42.6 ± 17.1% (mean ± SD, averaged over *N*_*r*_ = 4 biological replicates with a total of *N* = 426 cultures) of their surface area in the xy-plane due to cell disaggregation (**Supplementary Fig. 1b and d**). After this initial cell loss, most of the cell cultures recovered and grew until the end of differentiation (**Supplementary Fig. 1c**). The individual growth behavior of 3D cell cultures varied depending on the initial cell loss, but TF expression during differentiation was comparable for all cell cultures, where hiPSCs lost the expression of the pluripotency marker OCT4 within the first 48 h and gained the expression of the DE-specific markers, FOXA2 and SOX17 (**Fig. 1d**).

Furthermore, on-chip differentiation resulted in a nearly homogenous cell population with a yield as high as 96 ± 3% FOXA2/SOX17 double-expressing cells after 72 h (**Fig. 1e**, mean ± SD, *n* = 14936 nuclei in *N* = 20 cultures). However, experimental yields varied with an average of 90 ± 6% (mean ± SD, *N*_*r*_ = 4) of double-expressing cells at the end of differentiation, as previously observed in the standard cell well plate and chip culture formats ^15^.

We applied our platform to definitive endoderm (DE) differentiation, a critical first step for differentiating liver, gut, pancreas, lungs, trachea, and thyroid cell types. hiPSC-derived 3D cultures could be differentiated on-chip towards DE by activating the TGF-β/nodal and WNT signaling pathways with activin A and CHIR-99021, respectively. Over a period of 3 days, the two chemical components were controlled in time and concentration using fluid programming (**Methods**). Upon fixation of subsets of 3D cell cultures with NHS-ester every 24h, a differentiation trajectory was established (**Supplementary Fig. 2**). All cell cultures were immunostained on the chip at the end of differentiation for the pluripotency marker octamer-binding transcription factor 4 (OCT4) and the two DE-specific transcription factors (TFs) forkhead box A2 (FOXA2) and SRY-box 17 (SOX17). Bright-field and immunofluorescence (IF) images were recorded by standard confocal microscopy with a xy-resolution of 0.25 µm/px and a z-resolution of 1 µm/plane. Quantitative fluorescence signals of the TFs were extracted per nucleus for all 3D cell cultures by segmenting the 3D DAPI signal using a re-trained StarDist model ^13,14^. All IF images were corrected for the signal decrease in the z-direction caused by increasing light scattering and for the signal decrease in the xy-direction caused by gradually decreasing penetration of labeled antibodies (**Methods**).

### Label-free prediction of nuclei in 3D cell cultures

The full high-content imaging of hiSPC-derived 3D cell cultures with standard confocal microscopy with four fluorescence channels is laborious and time-consuming, requiring approximately 48 h for IF staining and 48 h of imaging. To cope with the chip throughput and characterize live cell cultures, we sought to predict the nuclear fluorescence of cells within whole 3D cell cultures along the DE differentiation trajectory from low-magnification bright-field images. For this purpose, we developed Bright2Nuc, a U-Net-based deep neural network (**Fig. 2a and b**). Bright2Nuc was trained on 255 paired bright-field stacks (one image below and one above a central slice) and fluorescence nucleus images of 181 live and 74 fixed 3D cell cultures by minimizing the mean-squared pixel error between the paired predicted and true IF nucleus images (see **Fig. 2a and Methods**). While images from fixed 3D cell cultures were acquired along the DE trajectory as described previously (**Fig. 1c**), images from live 3D cell cultures were obtained in the pluripotent state using a SOX2-T2A-tdTomato fluorescence reporter hiPSC line (**Supplementary Fig. 3**). Using the local 3D information from three neighboring z slices as input (**Fig. 2b**), the trained Bright2Nuc model accurately predicted nucleus images in an independent test set containing 85 cell cultures (26 fixed, 59 live) from the same experiments completely excluded from training (**Fig. 2c**).

**Figure 2.**
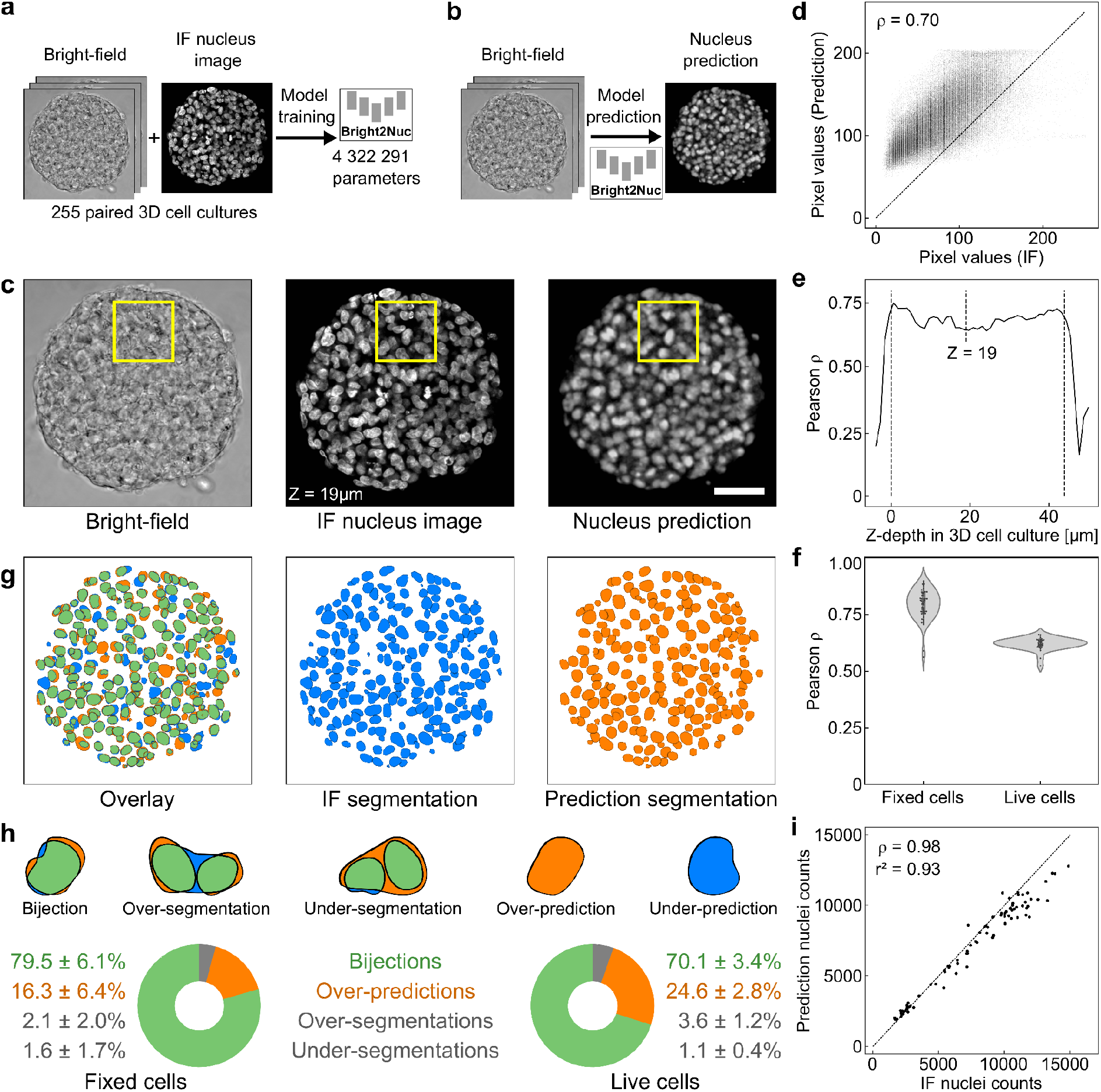
Neural network accurately predicts nuclear staining in 3D cell cultures from bright-field images. **a**, We trained Bright2Nuc, a U-Net-based neural network, on paired bright-field and IF nucleus images, and **b**, used it to predict nucleus images from bright-field images. **c**, Representative bright-field image of an on-chip 3D cell culture (left), corresponding ground-truth DAPI IF nucleus image (middle), and the predicted nucleus image (right). Yellow rectangles highlight the position of a specific region for comparison. Scale bar: 100 µm. **d**, Correlation between the fluorescence pixel values of the ground-truth nucleus image given in (**c**) and the pixel values of the corresponding predicted nucleus image (Pearson correlation *ρ* = 0.70). Each data point represents a single pixel value and the line represents the y = x identity. **e**, The model’s prediction performance is independent of the imaging depth. The Pearson correlation between the predicted and ground-truth pixels was stable along the z-axis of the cell culture, as shown in (c). **f**, Average Pearson correlation between the ground-truth and predicted pixel values from whole 3D cell cultures imaged in either fixed (*N* = 54) or live (*N* = 31) states. **g**, 3D segmentation of the IF nucleus image in (c) (middle, blue) and the predicted nucleus image (right, orange). The overlays of both segmentations show a high overlap (left, green; 80.6% overlapping area with the ground-truth segmentation). **h**, Top: Categories for comparison of predicted segmentation ground-truth IF segmentation. Bottom: Distribution of categories for predicted segmentations from fixed (*N* = 54) and live (*N* = 31) 3D cell cultures. **i**, High accuracy for the prediction of nuclei counts in the 85 fixed and live 3D cell cultures in the test set (Pearson correlation *ρ* = 0.98; coefficient of determination r^2^ = 0.93). The diagonal line represents the y = x identity.

The pixel values of the predicted nucleus images were highly correlated with the fluorescent nucleus signals of the ground-truth images (**Fig. 2d**, Pearson correlation of *ρ* = 0.70 for the pixel values of the image shown in Fig. 2c), independent of the tissue imaging depth (**Fig. 2e**). In the test set, predicted nucleus images from fixed 3D cell cultures showed an average pixel correlation of *ρ* = 0.79 ± 0.06 (mean ± SD, *N* = 54) that dropped to *ρ* = 0.62 ± 0.03 (*N* = 31) for live 3D cell cultures (**Fig. 2f**).

### Accurate segmentation of nuclei from bright-field imaging

The predicted nucleus images were segmented in 3D using a re-trained StarDist ^13,14^ model (**Fig. 2g**). To benchmark the results, segmented nuclei were classified into five categories: (i) bijections, that is, predicted nuclei overlapping with a single nucleus in the ground-truth image; (ii) over-segmentations, that is, two or more predicted nuclei overlapped with a single nucleus in the ground-truth image; (iii) under-segmentations, that is, single predicted nuclei overlapped with two or more nuclei in the ground-truth image; (iv) over-predictions, where predicted nuclei overlapped with no nuclei in the ground-truth image; and (v) under-predictions, where nuclei in the ground-truth existed with no overlap in the prediction image (**Fig. 2h**). In fixed cultures, the Bright2Nuc and StarDist approaches resulted in 79.5 ± 6.1% of the predicted nuclei as bijections (mean ± SD, *n* = 47224). An additional 2.1 ± 2.0% and 1.6 ± 1.7% of the predicted nuclei were correct, but over- or under-segmentated, respectively. However, the approach yielded 16.3 ± 6.4% of over-predictions, whereas conversely, 16.1 ± 6.6% of all ground-truth segmentations were under-predicted (**Fig. 2h**). Furthermore, prediction was enabled in live cultures at the cost of a slightly decreased accuracy of 70.1 ± 3.4% of bijections (**Fig. 2h**). The number of nuclei in each 3D cell culture was predicted with a high correlation to the groundtruth (**Fig. 2i**, *ρ* = 0.98; r^2^ = 0.93).

### Label-free prediction of differentiation state

Next, we investigated whether morphological and other features derived from bright-field images were predictive of hiPSC differentiation. In previous studies, the morphology of stem cell nuclei was shown to be indicative of their differentiation state ^16–19^. Therefore, we first mapped the expression of the three TFs to a single normalized value: the differentiation label (DL). The DL was defined as the ratio between the average of the normalized expressions of e_FOXA2_ and e_SOX17_ of the differentiation markers FOXA2 and SOX17, and the sum of the normalized expression of e_OCT4_ of the pluripotency marker OCT4 and the differentiation markers:

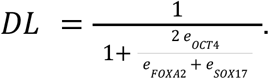

DL progressed from values close to zero for pluripotent cells at t = 0 h with high OCT4 expression to a value of ∼1 for DE cells after 72 h with high FOXA2 and SOX17 expression (**Supplementary Fig. 4**). To predict the DL for each nucleus, we designed an explainable feature-based approach. For each of the 26213 segmented nuclei along the DE differentiation trajectory, 120 features were extracted: 64 morphological features from the 3D segmentations after Bright2Nuc and StarDist applications and 56 bright-field texture features were extracted from a fixed-sized bounding box centered on the nucleus on the paired bright-field 3D images. A random forest algorithm with 1000 estimators and a depth of 10 was trained on the features for predicting DL (**Methods**). The ground truth DL for model training was calculated for each nucleus from the TF expression in the bounding box of the associated IF image (**Fig. 3a**).

**Figure 3.**
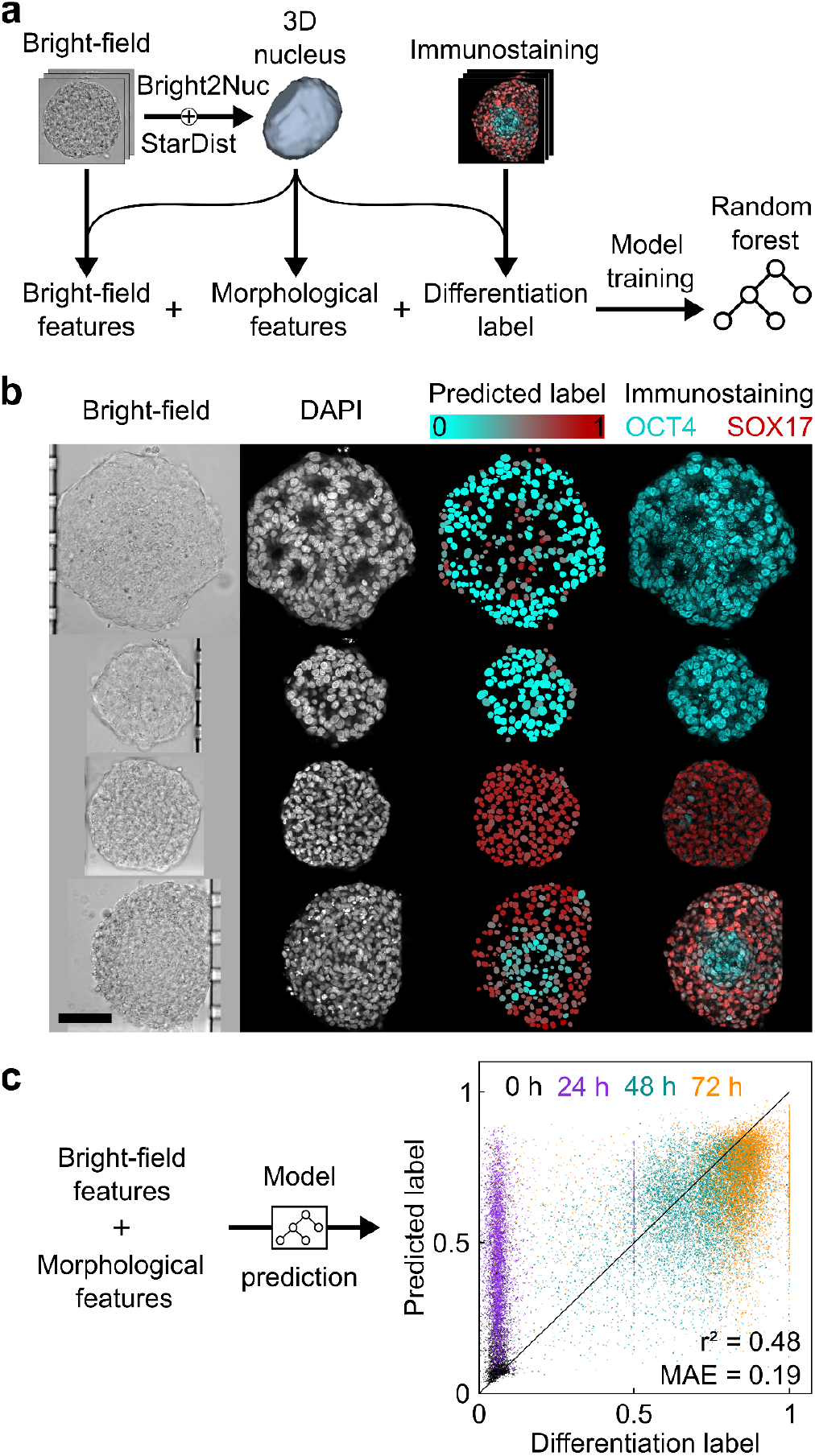
Bright-field and predicted morphological features predict single-cell differentiation state. **a**, We extracted 56 features from bright-field and 63 morphological features from segmented Bright2Nuc 3D images in combination with the differentiation label obtained from the IF expression to train the random forest model. **b**, Visually, the predicted label (3rd column, color coded from cyan to red) matches the true IF expression (last column, merged IF images showing OCT4 in cyan and SOX17 in red). Scale bar: 100 µm. The random forest model predicted a differentiation label for each cell in the on-chip 3D cell cultures. **c**, Comparison of the ground truth to the predicted differentiation label is shown on the right. Each data point represents a single nucleus (*n* = 26213) extracted from 49 3D cell cultures. (r^2^ = 0.48, mean absolute error = 0.19). The mean absolute error observed on nuclei in 3D cell cultures fixed 24 h after the start of DE differentiation is much higher (MAE_24h_ = 0.38 ± 0.18, *n* = 4376) than the one for other time points (MAE_0h_ = 0.15 ± 0.18, *n* = 3028; MAE_48h_ = 0.13 ± 0.11, *n* = 11364; MAE_72h_ = 0.17 ± 0.14, *n* = 7445). This discrepancy can be explained by the inaccurate value of the differentiation label obtained from the IF images. At 24 h, cells displayed high OCT4 and low FOXA2/SOX17 expression, thus resulting in low differentiation label values, even though the cells were already committed to the DE differentiation path. The random forest interestingly predicts a label value of DL_24h_-pred = 0.46 ± 0.18, which spans the range between the values for the 0 h and 48 h time points (DL_0h_-pred = 0.21 ± 0.18; DL_48h_-pred = 0.64 ± 0.15).

The resulting random forest model predicted single-cell DL in the bright-field images of 3D cell cultures fixed along the DE differentiation trajectory. In a round-robin fashion, five random forest models were trained on different training and test sets, ensuring that each cell culture was contained in the test set exactly once, while the training and test sets were split on the level of whole cell cultures. Visually, the predicted DL correlated with true IF staining, even in the case of non-homogeneously expressing cell cultures (**Fig. 3b**). For all 26213 nuclei, the predicted DL correlated with the true DL with a coefficient of determination of r^2^ =

0.48 and a mean absolute error of 0.19 ± 0.17 (mean ± SD, **Fig. 3c**). We observed strong deviations for nuclei fixed 24 h after DE induction, where the mean absolute error was higher than that at other time points. Interestingly, the predicted DL values at 24 h spanned the entire range, indicating that the random forest predicted nuclei as being in a developmental transition state rather than resembling pure DE or pluripotent cell states. The discrepancy between the predicted and true DL can be explained by an inaccurate ground truth at 24 h, where cells exhibited a high OCT4 and a low FOXA2/SOX17 expression (**Fig.1d-e**), resulting in a low true DL (0.09 ± 0.14, mean ± SD, *n* = 4376) indicative of pluripotent cells, which does not reflect the state of those cells after 24 h of DE induction. In fact, a single-cell transcriptomic time trajectory of DE differentiation under comparable conditions ^20^ showed that after 24 h, the transcriptome of hiPSCs clustered differently from pluripotent stem cells (**Supplementary Fig. 5a**), while still expressing Oct4 at the mRNA level (**Supplementary Fig. 5c**). Consequently, the simplified view of cell type differentiation based on single-cell state fluorescence markers argues for descriptive states, whereas the neural network learning approach resolves a continuous cell type transition, as is seen with time-resolved single-cell transcriptomics. Our approach highlights the rich and hidden information sources of bright-field images and their potential to resolve the transcriptional states of human stem cells.

### Label-free tracking of single cells

Bright2Nuc can bridge the domain gap between fixed and living tissues by tracking nuclei in live 3D cell cultures. To this end, we first generated ground-truth tracking data with 3D cell cultures formed from a mixture of wild-type hiPSCs and the SOX2-T2A-tdTomato reporter line in a one-to-two ratio to generate heterogeneous mixtures of labeled and non-labeled nuclei. Bright-field and IF images were acquired every 7.5 min for 17 h, resulting in 136 frames. The trained Bright2Nuc was then applied to every frame of the bright-field image sequence to predict nuclei. Predicted and IF-stained nuclei were tracked using TrackMate ^21^. Expectedly, we detected roughly two thirds of the number of nuclei (**Fig. 4a**, blue curve) in the IF images as compared to nuclei from the predicted images (**Fig. 4a**, black curve). With increasing experimental time, the number of detected tracks increased due to cell proliferation in both curves, with the ratio between ground-truth and predicted tracks remaining constant at *n*_GT_/*n*_Pred_=0.58 ± 0.03 (**Fig. 4a**, mean ± SD, *n* = 136 frames). Assigning nuclei to tracks resulted in a more stable nuclei count than counting segmented nuclei alone (**Fig. 4a**; orange curve). Tracks in the ground truth are, on average, longer than tracks found in the predicted sequence (**Supplementary Fig. 6b**), which can be due to under-predictions of Bright2Nuc (**Fig. 2**), causing nuclei to be lost in a track.

**Figure 4.**
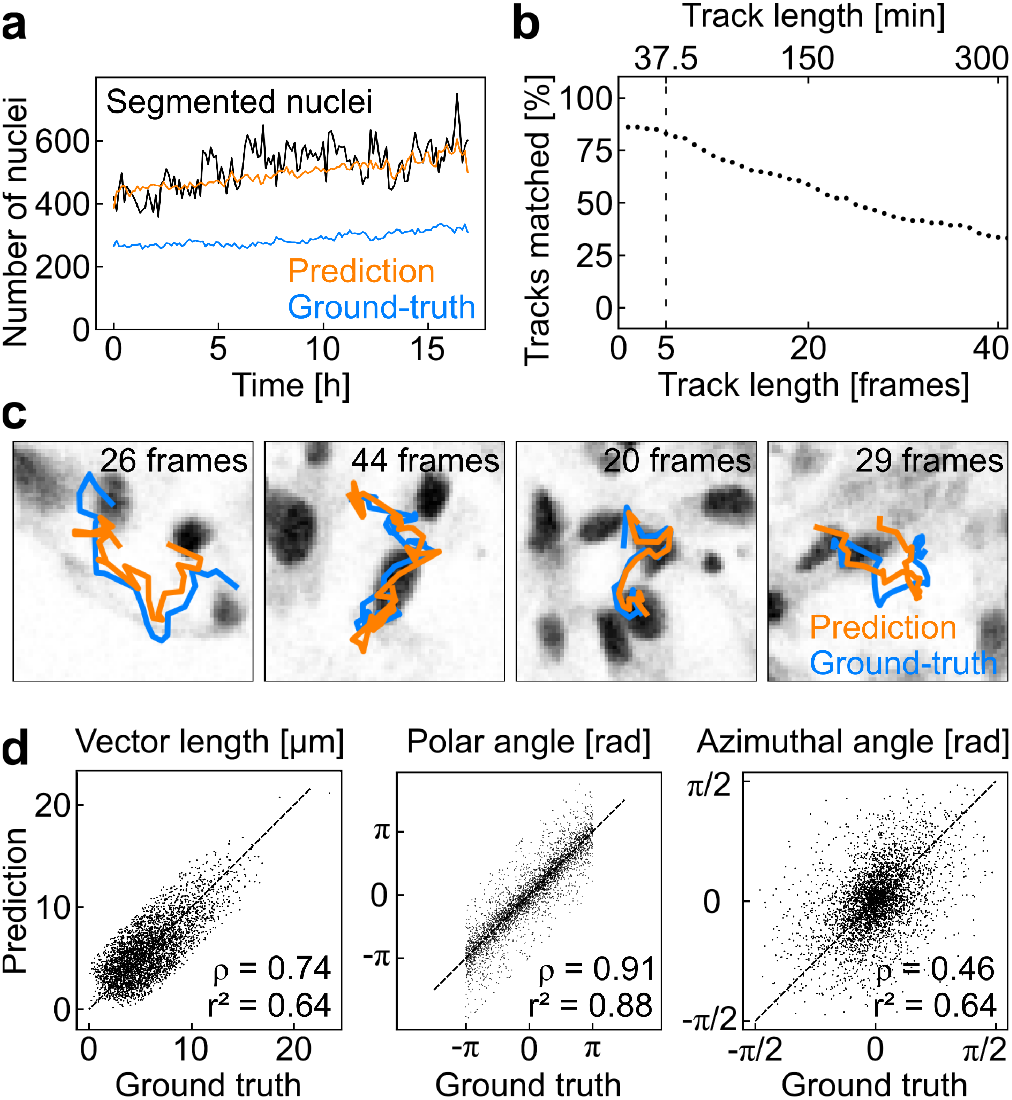
Label free single-cell tracking in live 3D cell cultures. **a**, Tracked nuclei in the label-free predicted images (orange) and fluorescence reporter images (blue) in one representative 3D cell culture. As per the ratio of fluorescent cells to the wild type, the ground-truth contained fewer tracks than the prediction. Incorporating temporal information by counting the number of tracks results in a more stable nucleus count than that based on the segmentations of each individual segmented label-free predicted image (black). **b**, If we require the tracks to match for a higher number of frames, the percentage of tracks in the fluorescence reporter images with a matched track in the predicted images decreases. The longer we require the tracks to be, the fewer we can identify them as matching. **c**, Representative 3D single-cell tracks in the predicted nucleus images (orange) and ground-truth SOX2-T2A-tdTomato fluorescence nucleus reporter images (blue). The background gray-value images show the ground-truth nuclear signal at the tracking end. The frame interval is 7.5 min. **d**, In spherical coordinates averaged over five frames, we found a high correlation in displacement (vector length, *ρ* = 0.74, r^2^ = 0.64) and directionality (polar angle, *ρ* = 0.91, r^2^ = 0.88, and azimuthal angle, *ρ* = 0.46, r^2^ = 0.64) and between prediction and ground truth.

For further tracking evaluation, we calculated the percentage of ground-truth tracks with a matched predicted track as a function of the predicted track length (**Fig. 4b**). We counted a match between two tracks if the distance between the center of mass of nuclei in the ground-truth and predicted images was less than 11 µm, which is roughly equal to the average nuclei diameter. The percentage of matched tracks decreased steadily from an initial 85% for the minimum requirement of two frames to approximately 50% for the requirement of 25 consecutive frames, corresponding to 3 h of imaging (**Fig. 4b**).

Finally, we compared tracks from the ground truth with their matched predicted tracks, taking an arbitrary threshold of five consecutive frames for which 81% of the ground-truth tracks had a predicted track match (**Fig. 4b**). Exemplary velocity vectors were derived by averaging the cell movement over five consecutive frames matched visually (**Fig. 4c**), and their spherical coordinates were correlated between the ground truth and predicted tracks with Pearson correlations for the vector lengths, polar angles, and azimuthal angles of *ρ* = 0.74, *ρ* = 0.91, and *ρ* = 0.46, respectively (**Fig. 4d)**. The distributions of both the ground truth and predicted velocity vector coordinates were similar (**Supplementary Fig. 6d-f**). Movement in the xy-plane was isotropic with a homogeneous distribution of the polar angle (**Supplementary Fig. 6e**), whereas the values of the azimuthal angle remained close to 0 (**Supplementary Fig. 6f**), indicating little movement in the z-direction. The limited z-directed movement was probably caused by the constrained geometry of the on-chip cultures. However, this confinement did not affect nuclei velocities, as evidenced by the lack of dependence of the velocity vector length on the z-position of the nuclei (**Supplementary Fig. 6c**). Overall, considering a track length of five frames, Bright2Nuc coupled with the TrackMate algorithm allowed us to track 81% of the nuclei in 3D cell culture (**Fig. 4c**).

### Label-free detection of cell velocities during differentiation

During DE differentiation, human pluripotent stem cells undergo synchronous epithelial-mesenchymal transition (EMT) ^22,23,24^. It is known that DE cells show significantly higher migration in 2D experiments as compared to pluripotent cells in a scratch assay ^23^, which argues that DE cells exhibit a higher cell mobility than stem cells. Therefore, in the last step, we investigated whether Bright2Nuc in combination with single-nuclei tracking can detect motility changes in label-free 3D cell cultures during DE differentiation. Along this line, we detected changes in the expression of cell adhesion molecules indicative of EMT during the DE differentiation on-chip (**Fig. 5a**). IF images of fixed 3D cell cultures along the DE differentiation trajectory showed that E-CAD/N-CAD expression changes co-occurred with upregulation of the DE marker SOX17. The single-cell transcriptomic dataset ^20^ also confirmed EMT TF changes, such as SNAIL1/2, whose expression peaked between the 24 h and 48 h time points (**Supplementary Fig. 5c**). To assess changes in cell motility during DE differentiation, we captured bright-field movies of six 3D cell cultures every 24 h over a period of 75 min with a 5 min/frame acquisition period (**Fig. 5**). Bright-field images were acquired using confocal microscopy with the same resolution as before. The Bright2Nuc + TrackMate approach was then used to extract the nuclei velocities, defined as the effective displacement of the nuclei over five frames (**Fig. 5a**). Strikingly, we observed an increase in effective cell displacement up to 48 h after induction of the DE differentiation. The highest effective cell displacement (D_48h_ = 0.98 ± 0.30 µm/min, mean ± SD) coincided with the time point of the E-CAD/N-CAD expression change and the appearance of the DE-specific marker SOX17. The population median of the four effective cell displacement distributions (D_0h_ = 0.78 ± 0.22 µm/min, D_24h_ = 0.70 ± 0.26 µm/min, D_48h_ = 0.98 ± 0.30 µm/min, D_72h_ = 0.82 ± 0.24 µm/min, mean ± SD) differs significantly (p = 3.9 × 10^−242^, Kruskal–Wallis H-test). Post-hoc analysis with a Bonferroni correction revealed a significant increase at the 48 h time point compared to the other time points (p_0h-48h_ = 7.3 × 10^−165^, p_24h-48h_ = 1.4 × 10^−164^, p_48h-72h_ = 2.3 × 10^−109^, Mann–Whitney-U-Test). Thus, the cells displayed higher motility during DE differentiation than during the pluripotent state. Notably, cell movements within the 3D cell cultures were non-directed throughout differentiation, as evidenced by the homogeneously low values of directionality calculated from the same velocity vectors (**Supplementary Fig. 7**). Interestingly, the measured increase in cell migration rates during DE differentiation was lower than that previously reported in 2D cultures ^23^, where DE cells showed a five times higher migration rate than that of pluripotent cells. This could reflect the more complex ECM within the 3D cell culture compared with a simple 2D cell surface culture.

**Figure 5.**
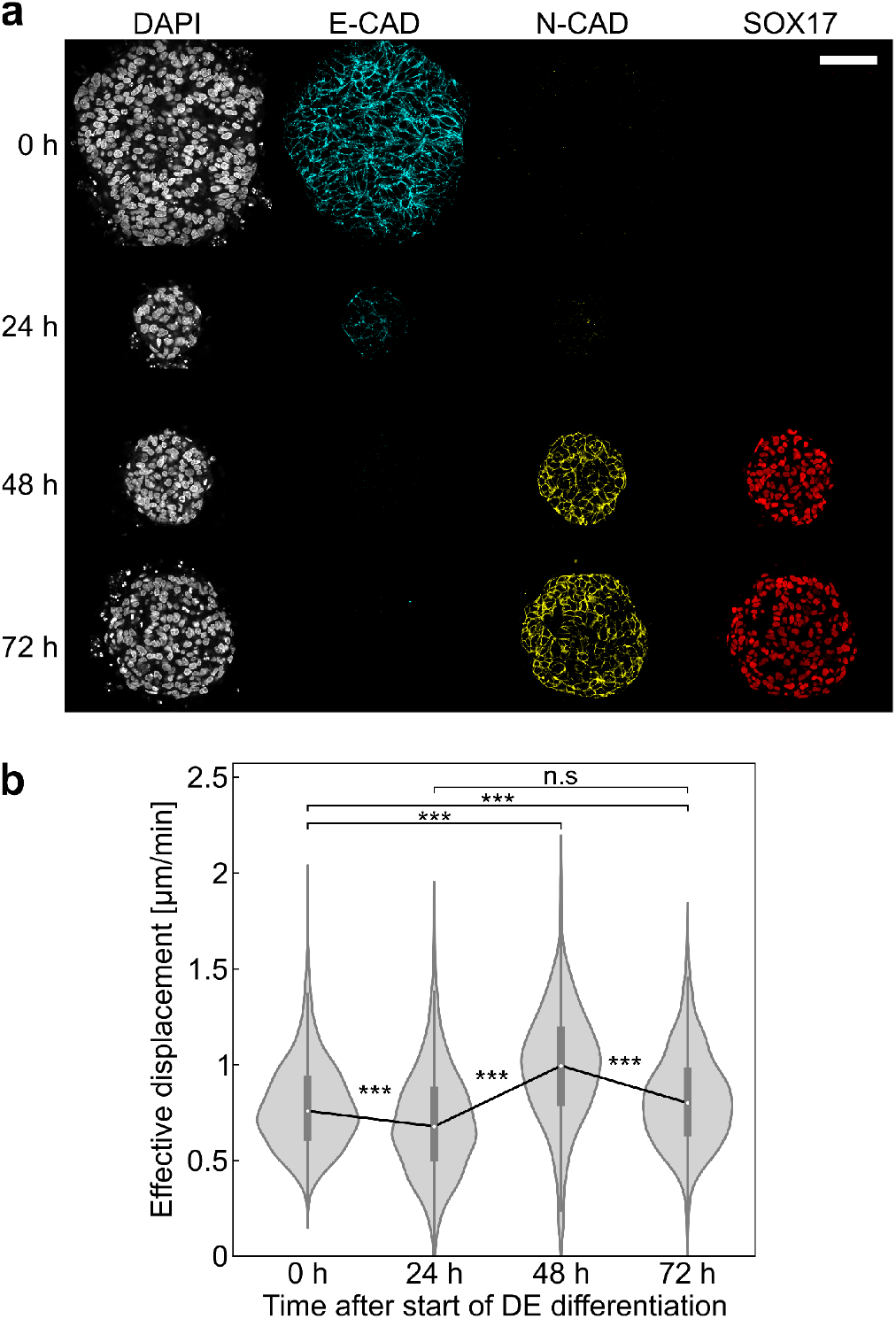
Single-cell live tracking in 3D cell cultures during DE differentiation captures increased cell mobility. **a**, Distribution of the effective displacement over five frames, calculated using Bright2Nuc and TrackMate on bright-field movies with a 24 h interval along the definitive endoderm differentiation trajectory. Each time point represents the distribution of the velocity vectors of nuclei in six 3D cell cultures (number of nuclei: *n*_0h_ = 2862, *n*_24h_ = 1308, *n*_48h_ = 2144, and *n*_72h_ = 2820). Movies were acquired with a 5 min per frame interval over 75 min. White circles indicate the mean, boxes mark the first and third quartiles, and bars indicate standard deviation. **b**, A significant increase in effective displacement correlates with the change in cadherin expression, as shown in the immunofluorescence confocal images of 3D cell cultures fixed along the DE differentiation trajectory, showing the evolution of epithelial-cadherin expression (E-CAD) and neural-cadherin expression (N-CAD). The evolution of the differentiation marker SOX17 was shown to mark progress towards the DE stage. Scale bar: 100 µm.

## Discussion

Microfluidic cell culture technologies for controlling the massive parallelization of 3D cell cultures are rapidly advancing, while high-throughput analytical methods to exploit miniaturized biological samples are lacking. In this study, we provide a label-free and live imaging approach to cope with the faster image acquisition of on-chip-cultivated 3D stem cell cultures. The developed neural network learning algorithm, Bright2Nuc, can be coupled with pre-existing image analysis tools such as StarDist and TrackMate for segmentation and tracking, respectively. With StarDist, Bright2Nuc enabled us to infer the cell number and nuclear locations from accessible confocal bright-field images with an accuracy of over 80% for 3D cell cultures with more than 10000 cells. Using the simplest imaging methods for high-content imaging, we offer a general imaging approach for screening 3D tissues. Deep learning approaches to detect cell nuclei have been previously employed for 2D adherent cell cultures ^8,9,11,12^ or histological tissue slices^10^. Existing neural networks have not yet attempted to infer information from cell images acquired with lower numerical aperture objectives and xy-resolution^9–11^ in 3D. Beyond locating only the nucleus volume and positions, morphological features from inferred nuclei and corresponding bright-field images allowed the determination of transcriptional cell states along the pluripotent to DE differentiation trajectory. Notably, the resulting cell state predictions were more continuous than the cell state descriptions generated from the fluorescence signals of descriptive fluorescence markers, which is the standard research approach in developmental biology. Our results highlight the currently untapped potential information contained in the bright-field images.

In addition to the static view, we demonstrated that Bright2Nuc can resolve real-time information from live nuclei in 3D cell cultures by using a neural network together with the TrackMate algorithm. For example, we captured label-free cell dynamics, indicating epithelial-mesenchymal transition on a timescale of hours. The homogeneous transition from pluripotent to DE cell stage within the on-chip cultivated 3D cell culture was reflected in the nuclei velocity changes during differentiation. Nevertheless, the provided image analysis method will allow the assessment of motion directionality, cell migration, and neighboring cell-cell contacts upon formation of architecturally complex structures into endodermal tissue, such as pancreatic or liver cell types. To further decrease the image acquisition time and increase the time resolution for subsequent 3D cell cultures, we expect that Bright2Nuc can be adopted for widefield bright-field images. In summary, we believe that the general framework of Bright2Nuc opens a wide field of applications for the high-content screening of 3D cell cultures on chips.

## Code availability

Bright2Nuc is available as a pip-installable package, together with commented analysis scripts at https://github.com/marrlab/Bright2Nuc.

## Data availability

The dataset is made available via Zenodo

## Author contributions

SA cultured, stained, and imaged the cell lines. DW implemented the code, trained the models, and performed the computational experiments. SA and DW evaluated the experiments, designed the figures, and wrote the manuscript with CM and MM. DW and SSB developed a Bright2Nuc Python package. CM and MM supervised this study.

## Acknowledgements

We thank Fabian Theis, Henrik Semb, Sophia Wagner, Melanie Schulz, Benedikt Mairhörmann, and Valerio Lupperger for their discussions and ideas.

---

**Supplementary Figure 1.**
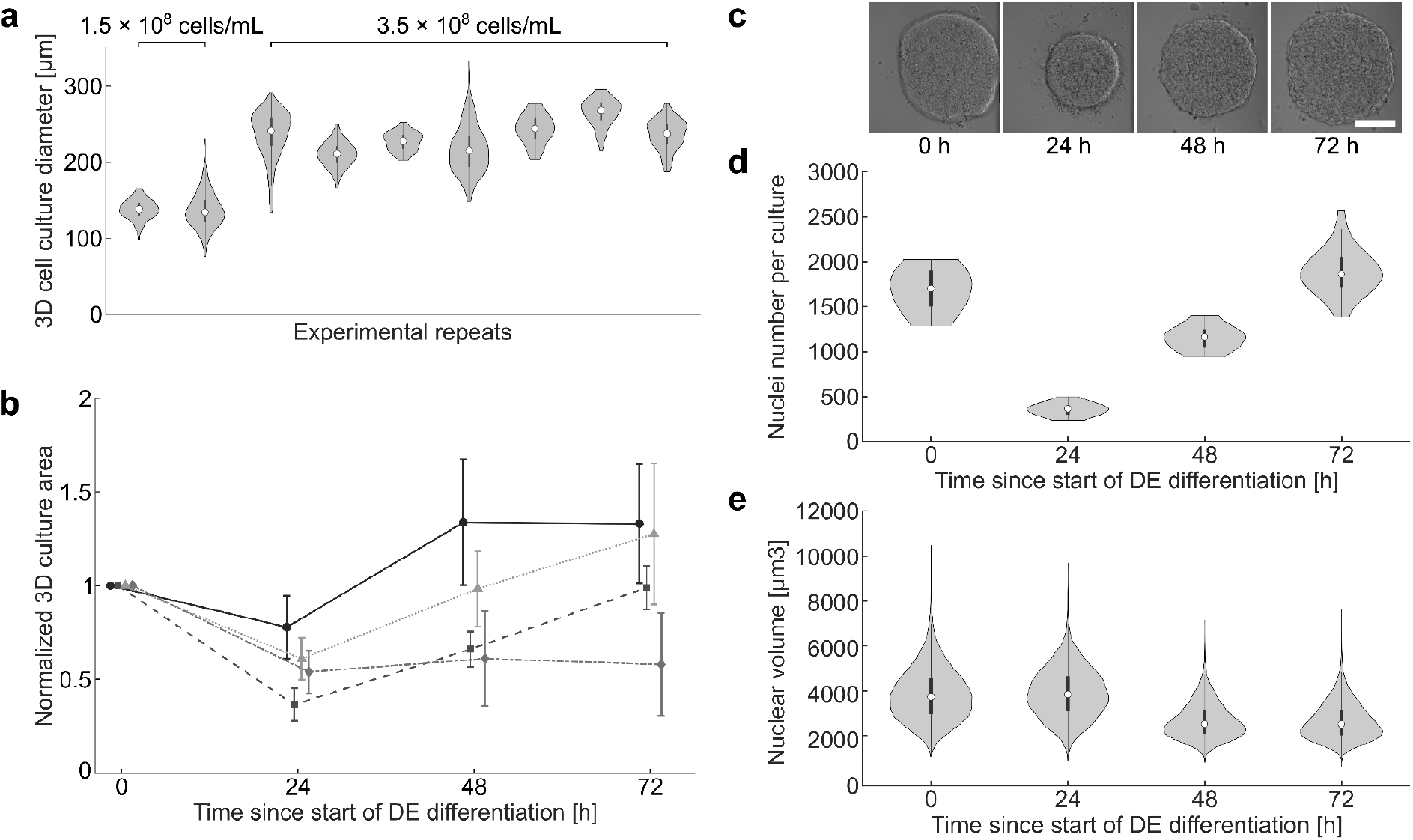
On-chip 3D cell culture and hiPSC differentiation. **a**, Distribution of 3D cell culture diameters measured 24 h after seeding cells on the microfluidic platform across nine experimental replicates. The first two repeats were seeded with a single-cell solution with a concentration of 1.5 × 10^8^ cells/mL; and the last seven repeats were seeded at a concentration of 3.5 × 10^8^ cells/mL. White circles indicate the mean, boxes mark the first and third quartiles, and bars indicate standard deviation. **b**, Evolution of the area occupied by 3D cell cultures measured in the xy-plane every 24 h along the DE differentiation trajectory. Cell culture areas were normalized individually to the area measured before the start of differentiation. Data points represent the mean and error bars show the standard deviation. Each of the four curves represents a distinct biological replicate with *N* = 128 cell cultures. **c**, Bright-field images of a typical 3D cell culture taken every 24 h along the DE differentiation trajectory. **d**, Evolution of the number of nuclei contained in 3D cell cultures fixed every 24 h along the DE differentiation trajectory for one experiment (curve with squares in **b**) with the corresponding evolution of the nuclear volume (**e**). White circles indicate the mean; boxes mark the first and third quartiles; bars indicate the standard deviation (*N*_0h_ = 16 cultures, *N*_24h_ = *N*_48h_ = *N*_72h_ = 36; *n*_0h_ = 26973 nuclei, *n*_24h_ = 12802, *n*_48h_ = 41550, and *n*_72h_ = 67780).

**Supplementary Figure 2.**
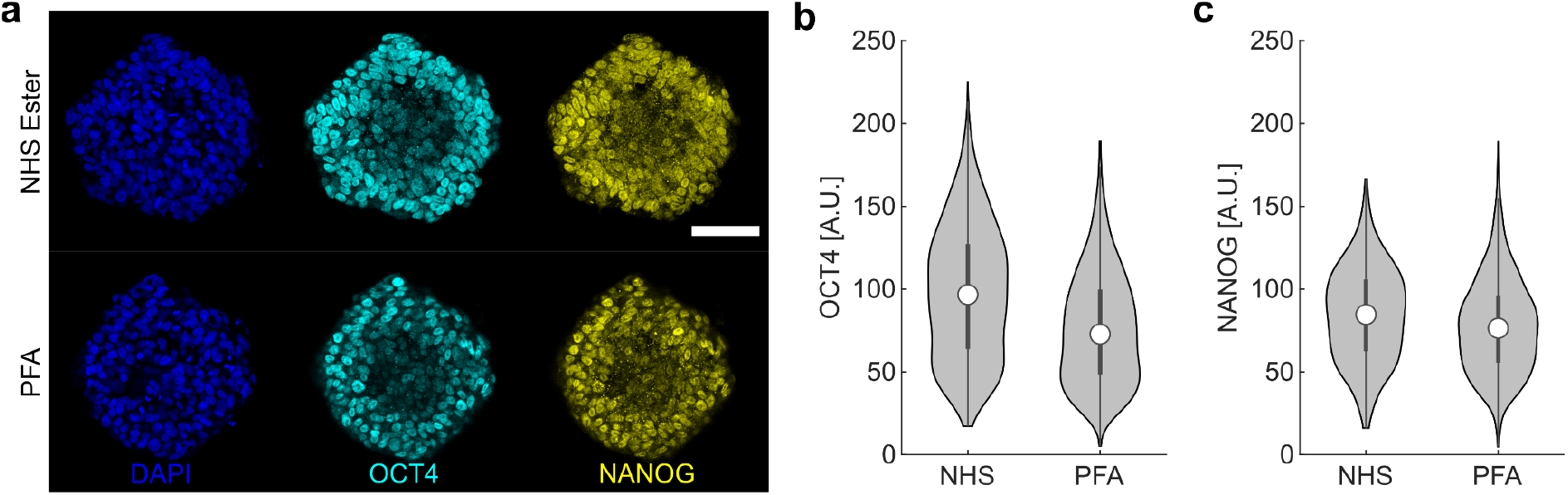
Comparison of fixation methods to enable time trajectories on the mLSI chip. **a**, Immunostaining images of iPSC 3D cell cultures stained for pluripotency markers OCT4 and NANOG. The top cell culture was fixed using 40 mM Bis-PEG4-NHS ester, while the bottom was fixed using 4% PFA. Scale bar: 100 µm. **b-c**, Mean fluorescence [A.U.] for OCT4 and NANOG in individual nuclei measured from confocal images after segmenting the DAPI signal. The distribution of nuclei fluorescence signals is shown for the 3D cell cultures fixed either with NHS (*n* = 1706 nuclei in eight cell cultures) or with PFA (*n* = 2852 nuclei in 10 cell cultures). White circles indicate the mean, boxes mark the first and third quartiles, and bars indicate standard deviation.

**Supplementary Figure 3.**
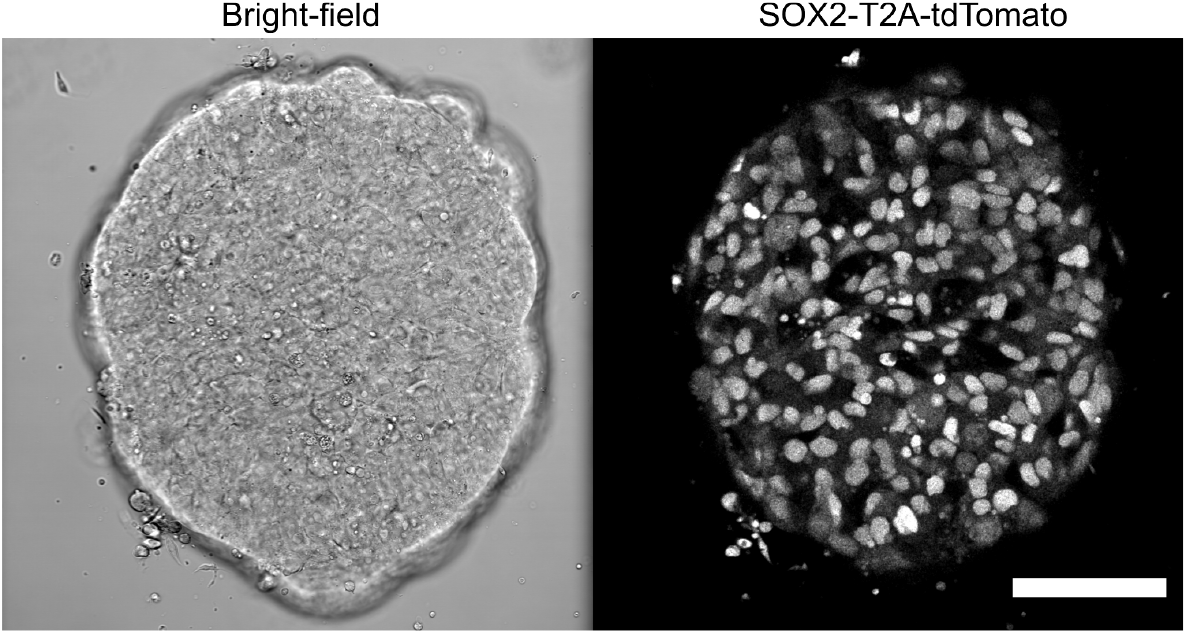
Confocal image of a pluripotent SOX2-T2A-tdTomato-reporter live 3D cell culture. Scale bar: 100 µm.

**Supplementary Figure 4.**
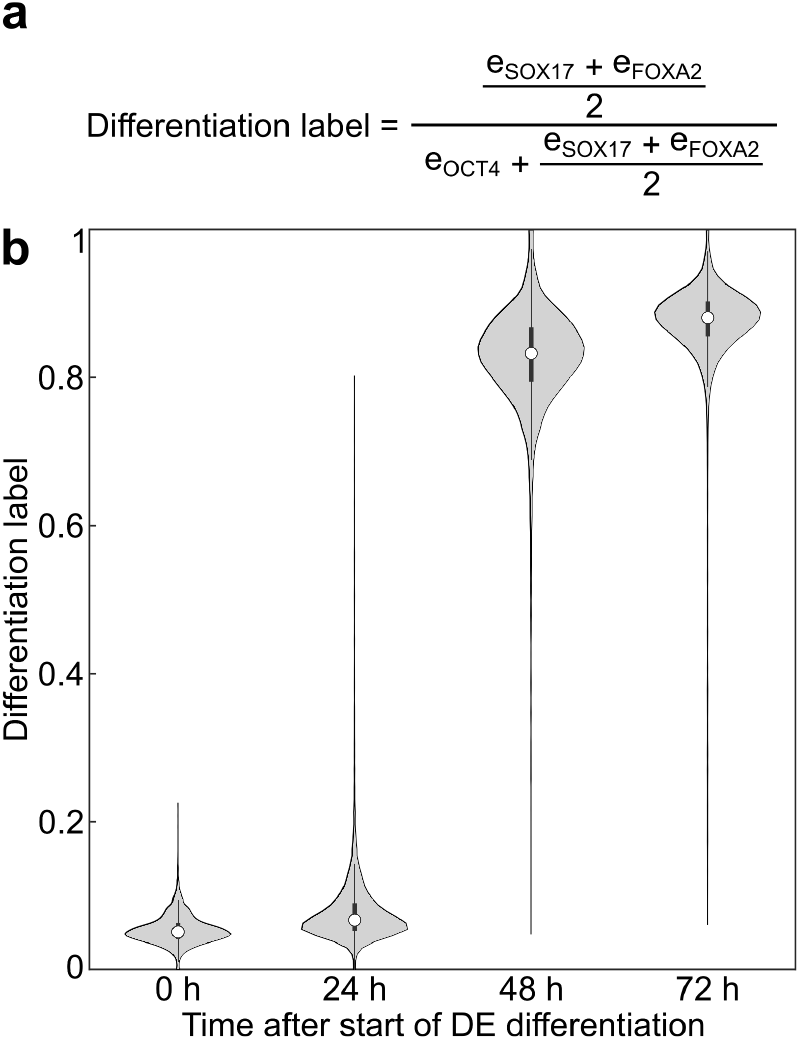
The differentiation state of cells expressed as differentiation label. **a**, The differentiation label is defined as the ratio of the average normalized expression of the differentiation markers e_FOXA2_ and e_SOX17_ to the sum of pluripotency (e_OCT4_) and differentiation expression. **b**, Evolution of the differentiation label along the definitive endoderm differentiation trajectory for the experiment presented in Fig. 1 (nuclei considered: *n*_0h_ = 5501, *n*_24h_ = 8700, *n*_48h_ = 22587, *n*_72h_ = 14936; 3D cell cultures: *N*_0h_ = 12, *N*_24h_ = 25, *N*_48h_ = 24, and *N*_72h_ = 30). White circles indicate the mean, boxes mark the first and third quartiles, and bars indicate standard deviation.

**Supplementary Figure 5.**
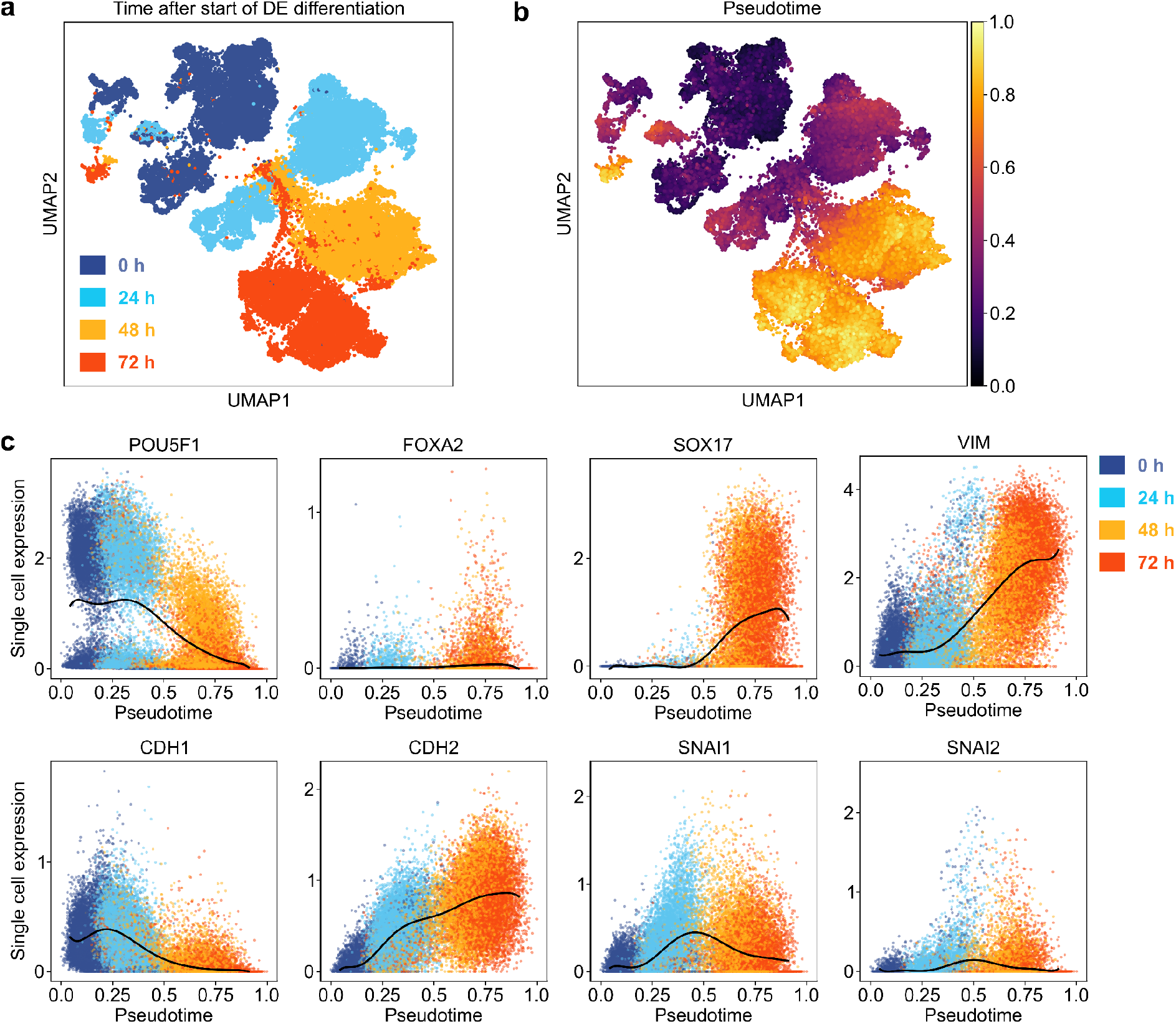
Single-cell transcriptomics of hiPSCs during definitive endoderm differentiation. Data adapted from datasets published by Cuomo et al. ^20^ **a**, UMAP clustering of single-cell RNA expression of human induced pluripotent stem cells sampled every 24 h during a 72 h differentiation protocol towards definitive endoderm. The overlaid colors represent the sampling time after the start of the DE differentiation. **b**, UMAP clustering overlaid with pseudotime, a measure of the differentiation state of the cells towards DE, analogous to our differentiation label. **c**, Single cell expression of selected genes as a function of pseudotime; colors indicate the sampling time. POU5F1, CDH1, and CDH2 are known as OCT4, E-CAD, and N-CAD, respectively. VIM is a mesenchymal marker that typically appears after the EMT. SNAI1 and SNAI2 are markers closely associated with EMT; interestingly, their expression peaks exactly between 24 h and 48 h.

**Supplementary Figure 6.**
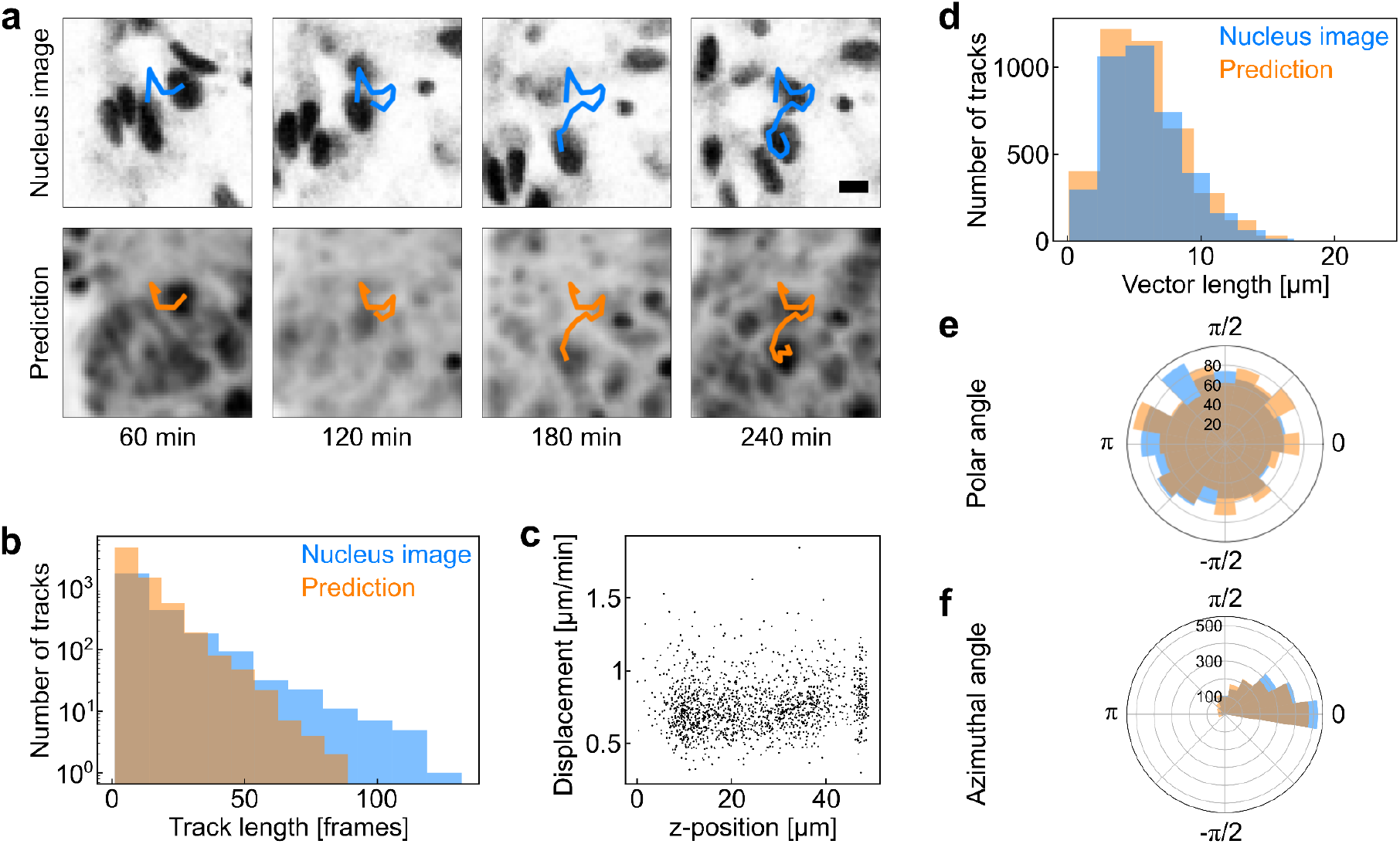
Single-cell live tracking in 3D cell cultures. **a**, Example frames with a 60 min interval for the tracking of a nucleus using both the ground-truth nucleus image (top row, SOX2 expression of pluripotent iPSCs) and *in silico* prediction from Bright2Nuc (bottom row). The overlaid tracks can be seen to have similar shapes in the ground truth and prediction. Scale bar: 5 µm. **b**, Distribution of track lengths. More vectors with a shorter length were found within the Bright2Nuc image frames than for the nucleus image tracks, as the tracking has a longer temporal persistence in the ground truth. **c**, Displacement of individual nuclei as a function of their z-position in the 3D cell culture. There was no observed dependence on the displacement relative to the z-position. **d-f**, Distribution of velocity vector coordinates for vectors with a mean error smaller than 11 µm between the positions of the nuclei in the prediction and the nuclear signal. Both distributions were remarkably similar in length (d), polar angle (e), and azimuthal angle (f). The mean velocity vector was calculated for five consecutive frames for 5362 corresponding nuclei.

**Supplementary Figure 7.**
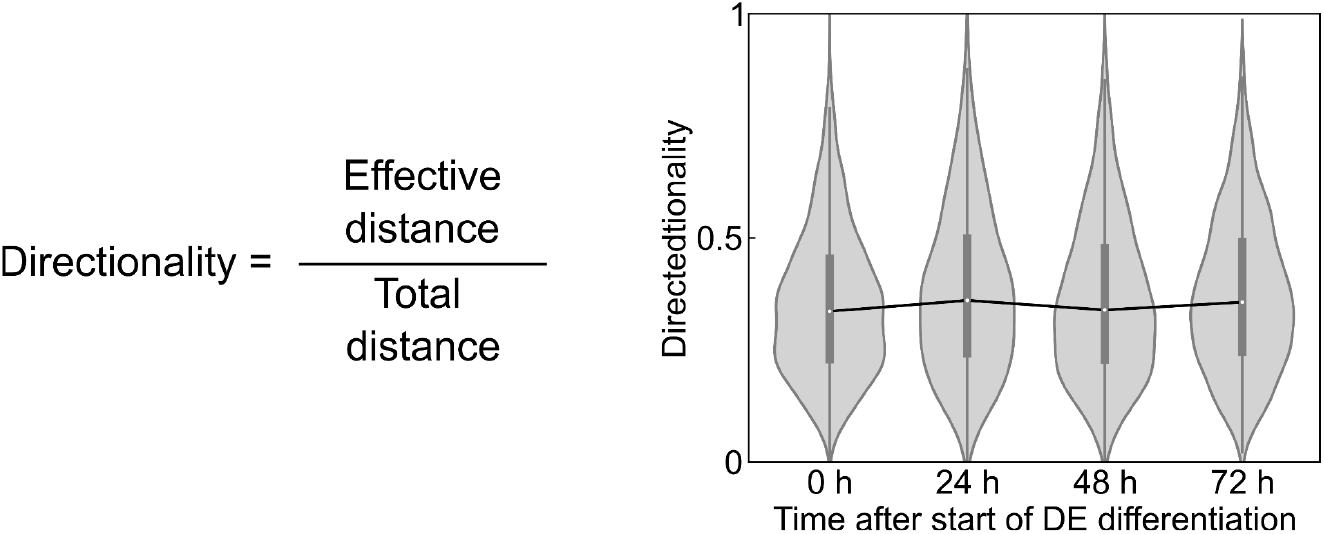
Directionality of cell movement in live tracking of 3D cell cultures during DE differentiation. Directionality, defined over five frames as the effective displacement divided by the absolute value of the displacement, is a measure of the randomness of cell movement. The directionality is calculated using Bright2Nuc and TrackMate on bright-field movies from Fig. 5 with a 24 h interval along the definitive endoderm differentiation trajectory. Each time point represents the distribution of the velocity vectors of nuclei in six 3D cell cultures (*n*_0h_ = 2862, *n*_24h_ = 1308, *n*_48h_ = 2144, and *n*_72h_ = 2820). Movies were acquired for 5 min. frame intervals over 75 min. White circles indicate the mean, boxes mark the first and third quartiles, and bars indicate standard deviation.

**Supplementary Figure 8.**
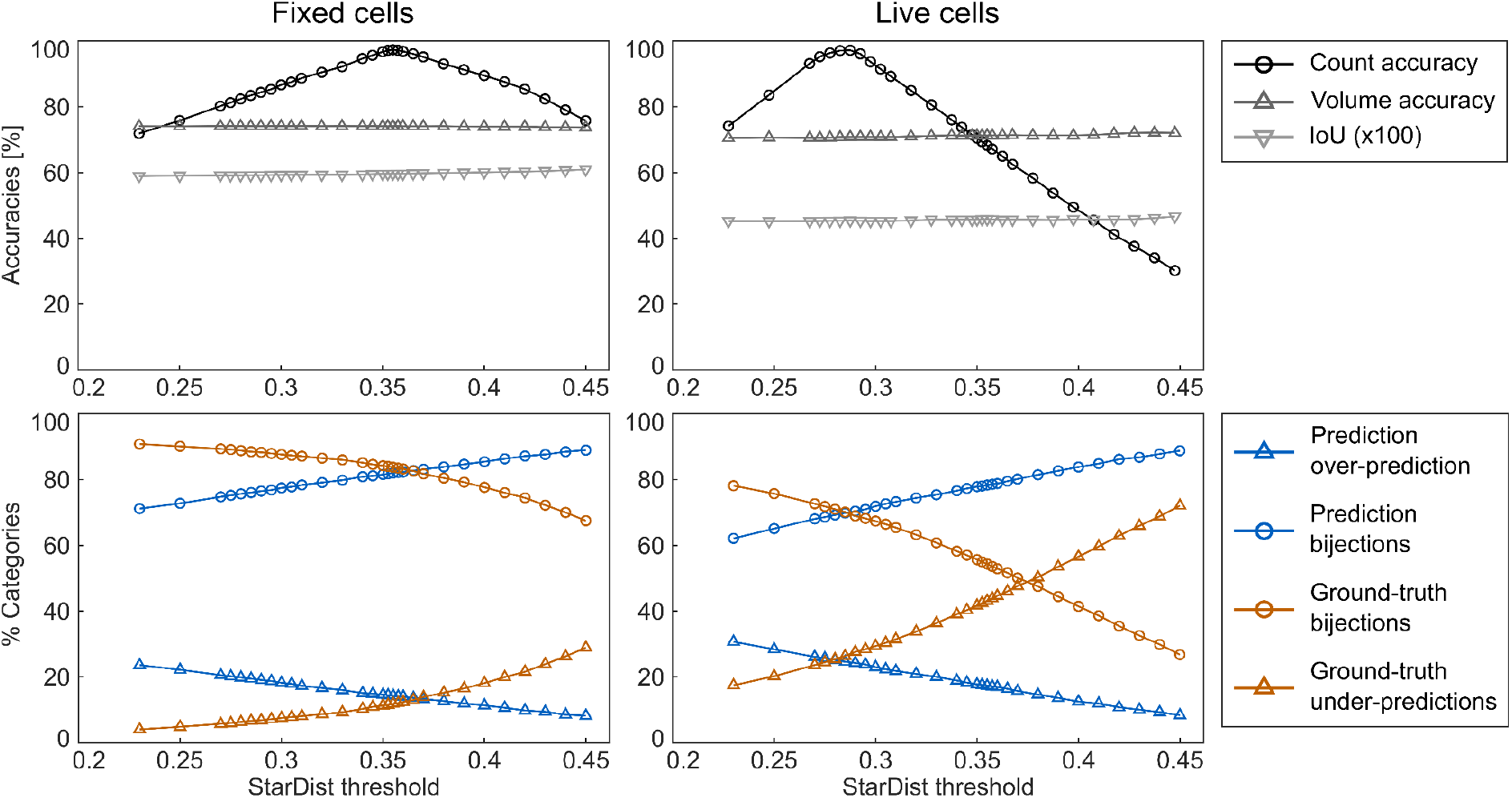
Threshold optimization for StarDist segmentation. The probability threshold, inherent to the StarDist model, for the segmentation of the *in silico* prediction nucleus image was optimized by maximizing the nuclei count accuracy calculated by comparing the number of objects segmented in the prediction with the number of objects in the ground-truth segmentation. Objects determined by the StarDist algorithm are accepted as valid segmentations if they have a probability that is higher than this threshold. Consequently, a higher threshold indicates fewer objects, but these objects are generally more trustworthy. Indeed, as the threshold increases, the percentage of over-predicted objects diminishes, and bijections increase as a percentage of *in silico* predictions. However, as fewer objects are predicted, the percentage of under-predictions increases as the bijections decrease as a percentage of ground truth objects. The value of the threshold has little to no consequence on the mean volume accuracy of the predicted bijections or their intersection over union ratio (IoU). It should be noted that, as a result, different threshold values were used when segmenting *in silico* prediction nucleus images of fixed cell cultures (threshold = 0.355) and live cell cultures (threshold = 0.290).

**Supplementary Video 1 | Seeding process of the microfluidic platform**.

Seeding of single-cell suspensions in the chambers of the microfluidic large-scale integration chip platform. Cells were perfused into each of the individually addressable cell culture chambers while the pneumatic membrane valves were in the open state. Cells in the four central cell compartments were shielded from direct flow upon the actuation of the pneumatic membrane valves. Subsequently, side channels were rinsed and the cell culture process, including a feeding cycle of every 4 h for 30 s, was started. Video dimension: 5570 × 3324 µm^2^. The video speed was real-time.

**Supplementary Video 2 | Live tracking of nuclei in pluripotent SOX2-T2A-tdTomato-reporter 3D cell culture**.

Video of a pluripotent SOX2-T2A-tdTomato-reporter 3D cell culture imaged with a confocal microscope with a 20X/0.8NA objective, as shown in Fig.4. Cell culture was imaged every 7.5 min for 17 h. (Left) Bright-field image of the 3D culture at a height of 5 µm. (Right) Average RFP fluorescence signal in the axial direction of the first 10 µm of the 3D cell culture. The nuclei in this video were tracked in 2D using StarDist integrated with TrackMate in ImageJ. The tracks are color-coded, and the lines indicate the past trajectories of the nuclei. Video height: 350 µm. Video speed:8 frames/s i.e. 3600×.

## Methods

### Microfluidic chip fabrication

PDMS chips were produced using traditional soft lithography techniques for two-layered devices ^25^. Briefly, two molds were fabricated by photolithography. The mold for the control channel network was made to have features 25µm in height using SU-8 3025 (MicroChem). The mold for the flow flayer was fabricated in three steps. First, the flow channel was made using the AZ40XT photoresist (MicroChemicals) at a height of 40µm and re-flowed to obtain rounded half-channels. Second, the perfusion channels of the cell culture chambers were produced at a height of 20µm using SU-8 3025 (MicroChem). Finally, the cell chambers were made using SU-8 3050 at a height of 50µm. All masks for photolithography were designed using the AutoCAD software (AutoDesk, 2019) and photoresists were patterned using a laser Micro Pattern Generator (µPG101, Heidelberg Instruments). All molds were coated with CYTOP™ (CTL-809M; AGC Chemicals) to prevent the adhesion of the PDMS. Chips were cast from the molds using Sylgard 184 PDMS (Dow Corning). The chips were assembled in a push-down configuration with the flow layer on the bottom. Flow and control layers were bonded using the off-ratio PDMS bonding procedure (5:1 pre-polymer to cross-linking reagents ratio for the control layer and 20:1 ratio for the flow layer). Finally, the flow layer was bonded to a glass slide carrier (Brain Laboratories) through oxygen plasma activation (20 W at 0.9 mbar for 25 s). Each cell culture chamber comprised a central flow channel and two side channels bifurcating at the entry of the culture chamber. Pneumatic membrane valves divided the central flow channel into four 640×400µm^2^ cell culture compartments with a volume of 0.013 µL. Gaps with a cross-section of 20×20µm^2^ allowed the crossing of fluids between the side channels and the culture compartments.

### Microfluidic chip operations

Pneumatic microfluidic valves (PMVs) within the chips were operated by applying a 1.5 mbar pressure on the control lines. The pressure could be switched on and off automatically on each individual control line using a homemade system enabling the individual control of 24 solenoid valves (LMV155RHY-5A-Q; SMC). Additionally, the system included a pressure regulator (ref pressure regulator) connected to 8 additional solenoid valves which applied a controllable pressure (0-1.5 bar) on light-proof gas-tight bottles containing the reagents to be introduced in the chip. All connections were made using Tygon tubings (ND 100-80; Proliquid, Germany). Chips were placed in a microscope stage top incubator (STX, Tokai Hit®) for maintaining a 37°C and 5% CO_2_ humidified environment while allowing live imaging. Prior to cell seeding, the cell culture chambers were coated with a 10% Pluronic® F-127 (Cat#9003-11-6; Sigma-Aldrich) in phosphate-buffered saline (PBS) overnight to prevent cell adhesion to the glass substrate or PDMS walls. Adherent cells were harvested at 70-80% confluence with TrypLE Express (Cat# 12604013; Gibco) and resuspended in 40-60 µL at a 1.5-3.5 × 10^8^ cells/mL in maintenance medium (mTeSR™ Plus; StemCell Technologies) supplemented with 10µM ROCK inhibitor (Y-27632, Cat# sc-281642A; Santa Cruz Biotechnology). For the formation of 3D cell cultures, a homogeneous single-cell solution was introduced into each cell culture chamber with PMVs in the open state. Upon actuation of the PMVs in the cell culture chamber, cells in the four compartments were isolated from the main flow stream, and cells in the side channels could be rinsed. Once seeded, cells were left to self-aggregate for 4 h before the first media renewal; media was then renewed in the cell culture chambers every 2 h using a 100 mbar forward fluidic pressure. The average diameter of on-chip 3D cell cultures depended on the single-cell solution density, i.e., 138 ± 13 µm for a cell density of 1.5 × 10^8^ cells/mL and 243 ± 19µm for a cell density of 3.5 × 10^8^ cells/mL. The 3D cell culture formation process after seeding was robust, as demonstrated by a low average coefficient of variation CV = 0.10 ± 0.05 and a chip-to-chip variation as low as CV = 0.08 (**Supplementary Fig. 1**).

### Human induced pluripotent stem cells (hiPSC) culture

When not specified otherwise, experiments were conducted using a hiPSC cell line ^26^ which was kindly provided by Alexander Kleger from Internal Medicine I, University Hospital, Ulm, Germany. In some experiments for the acquisition of live fluorescence nuclear signals, a SOX2-T2A-tdTomato reporter hiPSC line was used, generated by Shahryari et al ^27^. All cell lines were maintained in a pluripotent state as 2D adherent monolayers in conventional cell culture plates coated with Geltrex (Cat# A1413302; Life Technologies), fed daily using mTeSR™ Plus maintenance medium (StemCell Technologies, Canada) and maintained in a humidified atmosphere at 37°C and 5% CO_2_. After reaching 70–80% confluence, the cells were passaged with 0.5 mM EDTA (Cat# A4892; AppliChem) in PBS. To enhance cell viability after splitting, the maintenance medium was supplemented with 10 μM ROCK inhibitor (Y-27632, Cat# sc-281642A; Santa Cruz Biotechnology) for the following 24 h. Mycoplasma-free cell culture was regularly confirmed using a MycoAlert™ Plus Mycoplasma Detection Kit (Cat# LT07-703; Lonza).

### Definitive endoderm differentiation

Before DE induction, cells were fed for 24 h with maintenance medium and ROCK inhibitor and another 24 h with maintenance medium only. DE differentiation was induced 48 h after on-chip seeding, the start of differentiation has been called 0 h in this paper. We followed the differentiation protocol as published in previous literature ^28^, where the basal medium **(Supplementary Tables 1 and 2**) was supplemented with 100 ng/ml activin A (AA) (Cat# 120-14-300; Peprotech) and 3 µM CHIR-99021 (CHIR, GSK3β inhibitor, Cat# 24804-0004; Tebu-bio) on the first day of differentiation. Basal medium supplemented with 100 ng/ml AA and 0.3 µM CHIR was added on the second day, and basal medium with 100 ng/ml AA was added on the third day of differentiation.

### Immunocytochemistry on-chip

Partial fixations of 3D cell cultures (fixation of a subset of cell cultures in a single microfluidic platform, while keeping other cell cultures alive) were performed using NHS Ester (Bis-PEG4-NHS ester, CAT# BP-21602; BroadPharma). At the end of all experiments, before immunocytochemistry, all cell cultures were fixed in 4% paraformaldehyde in deionized water for 1 h (even if previously partially fixed NHS ester). Cell membranes were permeabilized with 0.2% Triton X-100 and 100 mM glycine in deionized water for 6 h and blocked in a blocking solution containing 3% donkey serum, 10% fetal calf serum, 0.1% Tween-20, and 0.1% bovine serum albumin (BSA) in PBS. Primary antibodies were diluted in the blocking solution and incubated in the cell culture chambers for 24 h before being rinsed away with PBS. Secondary antibodies and DAPI were then diluted also in the blocking solution and incubated for 24 h. All cell culture chambers were thoroughly rinsed with PBS prior to imaging. All steps were conducted on-chip and at room temperature. A full list of the antibodies used is given in **Supplementary Table 3**.

### Image acquisition

IF images and live bright-field images were acquired using a laser scanning confocal inverted microscope (Zeiss LSM 880 Airyscan) controlled by the ZEN Black version number software with a 20x/0.8-NA (numerical aperture) objective (Zeiss Plan-Apochromat 20x/0.8 M27), with up to five 8-bit or 16-bit data channels per image: transmitted light (bright-field using either the 633 nm or the 561 nm wavelength laser), DNA labeled with DAPI, and three types of antibodies distributed in the green (488 nm excitation), yellow (561 nm), and red (633 nm) channels (see **Supplementary Table 3** for antibody list). All images were acquired with a 0.25 × 0.25 µm^2^ pixel size with the number of pixels being adjusted so that the scanning area could fit the imaged object in the frame (typical scanning areas were around 200 × 200 µm^2^). Confocal stacks were acquired to image the full 50 µm z-depth of the 3D cell cultures with a z-resolution of 1 µm. For the imaging of fixed tissue, the pixel dwell time was kept in the 1 - 2 µs range, depending on scanning area size, with a two-fold line averaging. For live tissue, pixel dwell times were reduced to the 0.6 - 0.9 µs range, to accelerate imaging so that a full confocal stack of a 3D cell culture could be acquired in less than four minutes.

### Fluorescence signal correction

Prior to nuclear fluorescence signal quantification, two types of corrections were applied to the confocal image stacks. The first correction pertained to the weakening of fluorescence signals caused by the imaging depth in the 3D cell cultures. We calculated the relative average nuclear fluorescence signals (ratio between the average nuclear signal at a given depth and the average nuclear signal at the lowest point of the cell culture) as a function of the imaging depth. Under the assumption that nuclear signals should be constant on average relative to imaging depth, we calculated the imaging-depth decay rate through linear regression over all samples and for each wavelength. This wavelength-dependent correction was then applied to all image stacks. A second correction addressed antibody penetration inside the 3D cell culture during full-mount staining. Antibody penetration is a complex issue, which we resolved assuming that it factored in three main components: the antibodies, the x-y distance to the border of the cell culture, and the tissue type (differentiation state). We thus calculated the relative average nuclear fluorescence signal per antibody as a function of x-y distance to the border of the cell culture. The decay rates were calculated through linear regression per antibody for every sample within a time point to account for tissue type. All stacks were corrected under the assumption that nuclear signals should be constant on average relative to x-y penetration.

### In-silico nuclear staining

Bright2Nuc is a deep learning framework with a modified 3D U-Net ^29^, based on the InstantDL ^30^ package, designed to predict nuclear markers from bright-field images (**Fig. 2a-b**). Bright2Nuc works on 3D data, predicting the nuclear marker from slice to slice. To incorporate 3D information from the bright-field image stack, we added nearest neighbor slices to the input of the network. The U-Net architecture was modified from the 3D U-Net used in InstantDL by keeping the 3D convolutional layers the same but using anisotropic max-pooling in the encoder, and anisotropic upsampling in the decoder, keeping the z-dimension unchanged. Bright2Nuc can be trained and evaluated using 3D stacks, through which it will automatically iterate during training, Bright2Nuc will crop or pad the input data to a size of 384 pixels in x-y dimension. During inference, it can handle arbitrary image sizes. If desired, it can rescale the images to keep the nuclei diameter constant, which simplifies transfer learning between datasets. An average nucleus diameter of 30 pixels worked best for our model. All considered images were bilinearly downscaled by a factor of 2 to a 0.5 µm/px xy-resolution before being processed for further analysis. We trained Bright2Nuc (i) on 162 fixed 3D cell culture images in 3D in bright-field and with a DAPI staining and (ii) on 93 cell cultures recorded in 3D in live tissue in bright-field and with a SOX2-T2A-tdTomato reporter signal. Together, we used approximately a quarter million nuclei/cells. For training, we optimized a mean squared error loss for 50 epochs with a batch size of 5. For data augmentation, we used brightness and contrast shifts of 30% of the pixel values, horizontal and vertical flips, and zooms with a maximum of 30% of the image size. The test set contained 85 cell cultures split in a stratified manner from the experimental data, summing up to around 80.000 cells/nuclei. Bright2Nuc is a ready-to-use python package, it can be downloaded via GitHub or installed using pip install and comes with our pre-trained model.

### Nuclei segmentation

We trained two StarDist ^14^ 3D models, one to segment cell nuclei from images of a nuclear marker staining (stardist_nuc), and one to segment 3D cell cultures with an *in silico* staining (stardist_silico). We manually segmented five 3D cell cultures and trained stardist_nuc on the corresponding DAPI staining. To train stardist_silico, we created a training set of 15 3D cell cultures. We used the same five manually segmented cell cultures with the corresponding DAPI staining and the same five *in silico* stained cell cultures to the training set. Additionally, five *in silico* stained 3D cell cultures from the SOX2-T2A-tdTomato-reporter dataset were added, for which the segmentation ground truth was obtained by segmenting the SOX2-T2A-tdTomato signal using the first Stardist model trained on DAPI and manually verifying the results, summing up to 15 cell cultures. We trained the models as described in the StarDist documentation, but changed the anisotropy to two (x : 0.5, y : 0.5, z : 1) to match our imaging resolution. To adapt the StarDist model to the live tissue data, we found it sufficient to lower the StarDist threshold to 0.29 without retraining the stardist_silico model (**Supplementary Fig. 8**). With two resulting models (one for DAPI, one for in-silico) we segmented all DAPI stained 3D cell cultures and the *in silico* stained 3D cell cultures used in this paper.

### Transcription factor expression prediction

To assess the differentiation status of single nuclei, we formulated the expression of the pluripotent marker OCT4 (e_OCT4_), and the two differentiation markers FOXA2 (e_FOXA2_) and SOX17 (e_SOX17_) as a single ratio called differentiation label (DL) (**Supplementary Fig. 4**). Single-nuclei TFs expression were extracted from the corrected IF images (see Fluorescence signal correction in materials) by calculating the average fluorescence over each of the 3D DAPI StarDist segmentations. TFs expressions were normalized per dataset between 0 and 1 to the 1-th and 99-th percentiles, respectively. DL was then calculated for each nucleus as:

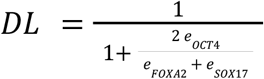

From each segmented nucleus, 56 bright-field and 64 morphological features were extracted. The data was then randomly split into five folds, with an 80% / 20% ratio between train and test set, and a random forest regressor was trained with 1000 estimators on each fold. In total, five random forest models were trained using the scikit-learn ^31^ framework in a round-robin fashion, splitting 20% of the nuclei into test sets iteratively, so that each nucleus was contained in the test set once.

### Live cell tracking

Single nuclei from the live *in silico* stained 3D cell cultures were tracked with Trackmate ^21^, using the integrated LoG detector with a sigma of 22 pixels (11 µm) and a quality threshold of 10. Data anisotropy was accounted for by setting the x- and y-values to 0.5 µm (leaving the z-value at 1 µm). The maximum linking distance was set to 22 pixels (11 µm) and the frame gap to two frames and 22 pixels. Tracking results were saved as a .csv file and evaluated in Python using pandas ^32^. Predicted tracks outside the cell culture were filtered out using neighborhood-based filtering, removing all tracks that have less than 5 neighbors in a distance of 50 pixels per time point. Effective displacements were calculated as the effective distance over five frames divided by the time, with the effective distance being the distance between the centers of mass of the considered nucleus between the first and fifth frames. Directionality was also calculated over five frames as the ratio of the effective distance and the total distance, with the total distance being the sum of distances between the centers of mass and between all considered frames.

---

**Supplementary Table 1.**
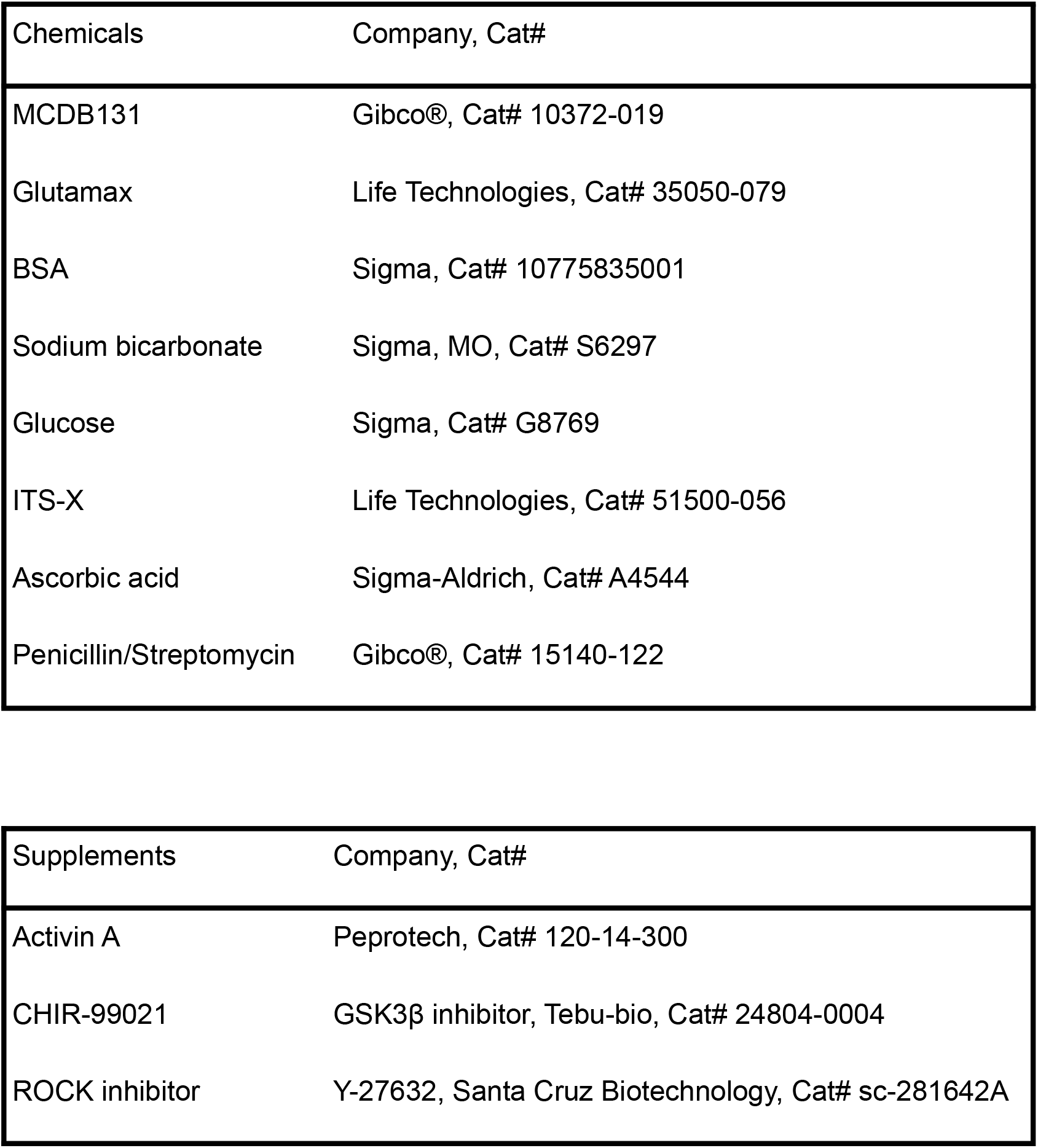
Chemical compounds used for DE differentiation.

**Supplementary Table 2.** Basal media composition and supplements used for DE differentiations

|  |
| --- |
| Basal media |
| MCDB131 |
| 1x Glutamax |
| 0.5% BSA |
| 1.5g/l Sodium Bicarbonate |
| 1.8g/l Glucose |
| 1% penicillin/streptomycin |

| Component | Day | Concentration |
| --- | --- | --- |
| ActA | Day01 | 100ng/ml |
| CHIR |  | 5μM |
| ROCK inhibitor |  | 0μM |
| ActA | Day02 | 100ng/ml |
| CHIR |  | 0.3μM |
| ActA | Day03 | 100ng/ml |
| CHIR |  | 0μM |

**Supplementary Table 3.** Antibodies used in immunochemistry

| Antibody | Host | Reference | Fluorophore |
| --- | --- | --- | --- |
| anti-OCT4 | Mouse IgG1 | Cell Signaling 75463 | N/A |
| anti-FOXA2 | Rabbit IgG | Cell Signaling 8186S | N/A |
| anti-SOX17 | Goat IgG | Acris/Novus GT15094 | N/A |
| anti-N-Cadherin | Mouse IgG1 | BD 610920 | N/A |
| anti-E-Cadherin | Rabbit IgG | Cell Signaling 3195S | N/A |
| anti-Mouse | Donkey | Invitrogen A21202 | Alexa Fluor 488 |
| anti-Rabbit | Donkey | Invitrogen A31572 | Alexa Fluor 555 |
| anti-Goat | Donkey | Invitrogen A21082 | Alexa Fluor 633 |
| anti-Mouse | Donkey | Dianova 107-175-151 | Alexa Fluor 647 |
| anti-Goat | Donkey | Invitrogen A11055 | Alexa Fluor 488 |

